# Historical biogeography of early diverging termite lineages (Isoptera: Teletisoptera)

**DOI:** 10.1101/2021.12.02.471008

**Authors:** Menglin Wang, Simon Hellemans, Jan Šobotník, Jigyasa Arora, Aleš Buček, David Sillam-Dussès, Crystal Clitheroe, Tomer Lu, Nathan Lo, Michael S. Engel, Yves Roisin, Theodore A. Evans, Thomas Bourguignon

## Abstract

Termites are social cockroaches distributed throughout warm temperate and tropical ecosystems. The ancestor of modern termites (crown-Isoptera) occurred during the earliest Cretaceous, approximately 140 million years ago, suggesting that both vicariance through continental drift and overseas dispersal may have shaped the distribution of early diverging termite lineages. We reconstruct the historical biogeography of three early diverging termite families – Stolotermitidae, Hodotermitidae, and Archotermopsidae – using the nuclear rRNA genes and mitochondrial genomes of 27 samples. Our analyses confirmed the monophyly of Stolotermitidae + Hodotermitidae + Archotermopsidae (clade Teletisoptera), with Stolotermitidae diverging from a monophyletic Hodotermitidae + Archotermopsidae approximately 100.3 Ma (94.3–110.4 Ma, 95% HPD), and with Archotermopsidae paraphyletic to a monophyletic Hodotermitidae. The Oriental *Archotermopsis* and the Nearctic *Zootermopsis* diverged 50.8 Ma (40.7–61.4 Ma, 95% HPD) before land connections between the Palearctic region and North America ceased to exist. The African *Hodotermes* + *Microhodotermes* diverged from *Anacanthotermes*, a genus found in Africa and Asia, 32.1 Ma (24.8–39.9 Ma, 95% HPD), and the most recent common ancestor of *Anacanthotermes* lived 10.7 Ma (7.3–14.3 Ma, 95% HPD), suggesting that *Anacanthotermes* dispersed to Asia using the land bridge connecting Africa and Eurasia ∼18–20 Ma. In contrast, the common ancestors of modern *Porotermes* and *Stolotermes* lived 20.2 Ma (15.7–25.1 Ma, 95% HPD) and 26.6 Ma (18.3–35.6 Ma, 95% HPD), respectively, indicating that the presence of these genera in South America, Africa, and Australia involved over-water dispersals. Our results suggest that early diverging termite lineages acquired their current distribution through a combination of over-water dispersals and dispersal via land bridges. We clarify the classification by resolving the paraphyly of Archotermopsidae, restricting the family to *Archotermopsis* and *Zootermopsis*, and elevating Hodotermopsinae (*Hodotermopsis*) as Hodotermopsidae (*status novum*).

## 1 Introduction

Termites are a clade of social cockroaches having a sister relationship with the wood-feeding cockroach genus *Cryptocercus* (Lo *et al*., 2000; Grimaldi & Engel 2005; Inward *et al*., 2007a, 2007b). The fossil record of termites dates back to the Early Cretaceous, ∼130 Ma (Thorne *et al*. 2000; Engel *et al*. 2016), and time-calibrated phylogenies suggest that the first termites appeared 140–150 million years ago (Ma) (Engel *et al*., 2009; Legendre *et al*., 2015; Bourguignon *et al*., 2015; Engel *et al*., 2016; Bucek *et al*., 2019). Therefore, the origin of termites predates the final stage of the breakup of Pangaea, and early diverging termite lineages may have a distribution based on vicariance through continental drift.

The first divergence amongst modern termites is that of Mastotermitidae and Euisoptera, the clade composed of all non-mastotermitid termites, ∼140–150 Ma (Inward *et al*., 2007a; Engel *et al*., 2009; Bourguignon *et al*., 2015; Bucek *et al*., 2019). The only extant species of Mastotermitidae, *Mastotermes darwiniensis*, is native to northern Australia and was introduced to New Guinea (Barrett 1965). However, fossils of *Mastotermes* have been unearthed in Russia, Mexico, the Dominican Republic, Brazil, Europe, Ethiopia, and Myanmar (Krishna & Emerson 1983; Krishna & Grimaldi 1991; Wappler & Engel 2006; Krishna *et al*., 2013; Vršanský & Aristov 2014; Engel *et al*., 2015; Zhao *et al*., 2019; Bezerra *et al*., 2020), indicating that *Mastotermes* was once globally distributed and acquired its modern relict distribution through multiple extinction events. The past global distribution of *Mastotermes* can be explained by a combination of vicariance through continental drift and transoceanic dispersal events. In addition, numerous extinct lineages of Mastotermitidae further attest to the greater past diversity of this lineage during the Cretaceous and Paleogene (Krishna *et al*., 2013). Unfortunately, molecular-based time-calibrated phylogenies cannot be used to resolve the historical biogeography of *Mastotermes*. However, the method can be used to resolve the origin and patterns of distribution of representatives of other early diverging termite families with broader extant diversity and occurrences.

The first divergence within the Euisoptera is the separation of Teletisoptera (Stolotermitidae + Hodotermitidae + Archotermopsidae) from Icoisoptera (Kalotermitidae + Neoisoptera), dated at 130–145 Ma (Bourguignon *et al*., 2015; Bucek *et al*., 2019). The most recent common ancestor of the former clade corresponds to the split between Stolotermitidae and Archotermopsidae + Hodotermitidae and was estimated at 80–115 Ma (Bourguignon *et al*., 2015; Bucek *et al*., 2019). Therefore, cladogenesis in Stolotermitidae + Hodotermitidae + Archotermopsidae was initiated before the final stage of the breakup of Pangaea, indicating that their current distribution may have been shaped by vicariance through continental drift (Bourguignon *et al*., 2015). Alternatively, despite their ancient origin, Stolotermitidae + Hodotermitidae + Archotermopsidae may have acquired their modern distribution by dispersal, with extensive extinction of stem-group Teletisoptera during the Cretaceous and perhaps early Paleogene. Indeed, the presence of several extinct groups that are putatively stem groups to this clade are known from the mid-Cretaceous (e.g., Arceotermitidae and Krishnatermitidae at 99 Ma: Jiang *et al*., 2021). A comprehensive phylogeny including samples collected across the range of these three early diverging termite families could help determine whether their modern distribution was shaped primarily by dispersal, vicariance, or a combination of these two phenomena.

Extant Stolotermitidae are found in Australia, South Africa, South America, and New Zealand, a distribution often interpreted as relict and reflecting an ancient occurrence across Gondwana prior to its initial breakup approximately 100 Ma (Krishna *et al*., 2013). Modern Hodotermitidae are distributed across the deserts of Africa, the Middle East, and South Asia. This distribution was possibly acquired as arid biomes gradually expanded during the Oligocene and Miocene (Edwards *et al*., 2010), following the global cooling that was initiated at the Eocene-Oligocene boundary, ∼34 Ma (Zachos *et al*., 2001). Finally, the Archotermopsidae have a disjunct distribution across the Northern Hemisphere, with *Archotermopsis* living at the foothills of the Himalayan region and in mountains of Vietnam; *Hodotermopsis* living in Vietnam, South China, and Japan; and *Zootermopsis* native to the western part of the Nearctic region (Krishna *et al*., 2013) and introduced to Japan (Yashiro *et al*., 2018). While the fossil record of the three families is more fragmentary than that of Mastotermitidae, most of these fossils indicate that the families once enjoyed a broader distribution. For example, the genus *Chilgatermes* from Oligocene deposits of Ethiopia is a relative of Porotermitinae (Stolotermitidae) (Engel *et al*., 2013), while *Termopsis* (of the extinct family Termopsidae, and likely related to Archotermopsidae + Hodotermitidae) is found in middle Eocene Baltic amber (Engel *et al*., 2007; Krishna *et al*., 2013). Similarly, the extinct archotermopsid genera *Ulmeriella* and *Gyatermes* are known from a variety of fossil deposits in Europe and Asia, as well as North America for the former (Engel & Gross 2009; Krishna *et al*., 2013; Engel & Tanaka 2015). Additionally, there are various extinct genera from the Cretaceous that are p utatively stem groups to the Teletisoptera, such as *Arceotermes* and *Cosmotermes* from the 99 Ma Kachin amber (Arceotermitidae: Jiang *et al*., 2021), *Cretatermes* from 95 Ma deposits in Labrador (Emerson 1967), and possibly *Carinatermes* in 90 Ma New Jersey amber (Krishna & Grimaldi 2000). Other fossils are summarized by Krishna *et al*. (2013) and Barden & Engel (2021). Thus, the historical biogeography of Teletisoptera may be more intricate than previously acknowledged.

The classification of these lineages has changed considerably over the last century (Table 1). At the time of Holmgren’s (1911) monumental classification of termites, he placed all of the aforementioned genera in his family ‘Protermitidae’. Therein he recognized several subfamilies, with the Hodotermitinae (*Archotermopsis, Hodotermes*), Stolotermitinae (*Stolotermes*), and Calotermitinae (including *Porotermes*) including those genera discussed herein. Most authors followed this system with subtle modifications, most noteworthy of which was the removal of kalotermitids to their own family. While maintaining a single family as Hodotermitidae, Emerson (1933, 1942) considered the genera *Hodotermopsis, Archotermopsis*, and *Zootermopsis* as related to the extinct genus *Termopsis* and placed them all within subfamily Termopsinae, relegating *Hodotermes, Microhodotermes*, and *Anacanthotermes* to Hodotermitinae, and *Stolotermes* and *Porotermes* were each placed in their own monogeneric subfamilies. Grassé (1949) was the first to emphasize the distinctive biology of these groups in the classification, elevating the so-called dampwood termites to family rank as Termopsidae and as more formally distinct from the harvesters of the Hodotermitidae s.str. (= Hodotermitinae sensu Emerson, 1942), a system that was maintained by most authors for the next 60 years (e.g. Engel & Krishna, 2004; Thorne *et al*., 2000; Weidner, 1955). The first major reconsideration of this system came with the morphological and paleontological phylogeny of Engel et al. (2009). In their analysis, *Termopsis* was recovered as unrelated to the modern genera of “Termopsidae”, necessitating the removal of the extant diversity to the Archotermopsidae, while most recently Jiang *et al*. (2021) separated *Hodotermopsis* into a monogeneric subfamily, Hodotermopsinae.

**Table 1.** Comparison of different classifications of extant basal Euisoptera. Fossil representatives are not covered here but are largely summarized by Krishna et al. (2013), Barden and Engel (2021), and Jiang *et al*. (2021). Families boldfaced in small caps, and genera color coded by clades.

|  | Emerson (1942),<br>Snyder (1949),<br>Krishna (1970) | Grassé (1949), Weidner<br>(1955), Engel &<br>Krishna (2004) | Engel *et al.* (2009,<br>2016), Krishna *et al.* (2013) | Jiang *et al.* (2021) | Herein |
| --- | --- | --- | --- | --- | --- |
| **PROTERMITIDAE**<sup>1</sup> | **HODOTERMITIDAE** | **HODOTERMITIDAE** | **HODOTERMITIDAE** | **HODOTERMITIDAE** | **HODOTERMITIDAE** |
| Hodotermitinae | Hodotermitinae | *Anacanthotermes* | *Anacanthotermes* | *Anacanthotermes* | *Anacanthotermes* |
| *Archotermopsis* | *Anacanthotermes* | *Microhodotermes* | *Microhodotermes* | *Microhodotermes* | *Microhodotermes* |
| *Hodotermes*<sup>2</sup> | *Microhodotermes* | *Hodotermes* | *Hodotermes* | *Hodotermes* | *Hodotermes* |
| Stolotermitinae | *Hodotermes* | **TERMOPSIDAE** | **ARCHOTERMOPSIDAE** | **ARCHOTERMOPSIDAE** | **HODOTERMOPSIDAE** |
| *Stolotermes* | Termopsinae | *Hodotermopsis* | *Hodotermopsis* | Hodotermopsinae | *Hodotermopsis* |
| Calotermitinae<sup>3</sup> | *Hodotermopsis* | *Archotermopsis* | *Archotermopsis* | *Hodotermopsis* | **ARCHOTERMOPSIDAE** |
| *Porotermes* | *Archotermopsis* | *Zootermopsis* | *Zootermopsis* | Archotermopsinae | *Archotermopsis* |
|  | *Zootermopsis* | Porotermitinae | **STOLOTERMITIDAE** | *Archotermopsis* | *Zootermopsis* |
|  | Porotermitinae | *Porotermes* | Porotermitinae | *Zootermopsis* | **STOLOTERMITIDAE** |
|  | *Porotermes* | Stolotermitinae | *Porotermes* | **STOLOTERMITIDAE** | Porotermitinae |
|  | Stolotermitinae | *Stolotermes* | Stolotermitinae | Porotermitinae | *Porotermes* |
|  | *Stolotermes* |  | *Stolotermes* | *Porotermes* | Stolotermitinae |
|  |  |  |  | Stolotermitinae | *Stolotermes* |
|  |  |  |  | *Stolotermes* |  |

<sup>1</sup> Holmgren's (1911) Protermitidae also included Mastotermitinae, not covered herein.
<sup>2</sup> Holmgren (1911) included *Anacanthotermes* as a subgenus of *Hodotermes*.
<sup>3</sup> Holmgren (1911) also included in this subfamily *Calotermes* (= *Kalotermes s.l.*, or what today is recognized as Kalotermitidae).

While the historical biogeography of Neoisoptera and Kalotermitidae has been studied in detail (Bourguignon *et al*., 2016, 2017; Wang *et al*., 2019; Romero Arias *et al*., 2021; Bucek *et al*., 2021), only a few species of Stolotermitidae, Hodotermitidae, and Archotermopsidae have been included in previous termite phylogenies. In this paper, we carried out a representative sampling of species belonging to these three families. We obtained the nuclear ribosomal RNA genes (5S, 5.8S, 18S, 28S) and mitochondrial genomes of 27 samples collected across the distribution of the group. We used this dataset to reconstruct time-calibrated phylogenies, clarify the classification, and shed light on the historical biogeography of these early diverging termite lineages.

## 2 Materials and Methods

### 2.1 Sampling and sequencing

We sequenced eight samples of Stolotermitidae, eight samples of Archotermopsidae, and seven samples of Hodotermitidae. In addition to these 23 samples, we also sequenced 36 termite species belonging to other families that we used as outgroups, including 17 species of Termitidae, 11 species of Rhinotermitidae, seven species of Kalotermitidae, and the only species of Mastotermitidae, *M. darwiniensis*. We combined these sequences with previously published mitochondrial genomes of one species of Stolotermitidae, two species of Archotermopsidae, one species of Hodotermitidae, *M. darwiniensis*, and one species of Cryptocercidae. Our final dataset comprised sequence data for 64 termite species and one non-termite cockroach species, *Cryptocercus punctulatus* (Table S1).

Whole genomic DNA was extracted with the DNeasy Blood & Tissue kit using complete individuals, including guts. The concentration of DNA was measured with Qubit 3.0 fluorometer and adjusted to a concentration of 0.5 ng/μl. The library of each sample was prepared separately with the NEBNext® Ultra (tm) II FS DNA Library Preparation Kit and the Unique Dual Indexing kit (New England Biolabs), with reagent volumes reduced to one-fifteenth of that advised by the manufacturer. We retained the enzymatic fragmentation step during library preparation for the few samples collected for genomic analyzes and preserved in RNA-later® at -20°C or -80°C until DNA extraction. However, most samples were collected over the past decades in alcohol and stored at room temperature for taxonomic purposes. Because the DNA of these samples was typically highly fragmented, we prepared libraries without the enzymatic fragmentation step. Libraries were pooled together and paired-end sequenced with the Illumina sequencing platform at a read length of 150 bp.

### 2.2 Assembly and Alignment

We checked read quality using Fastp v0.20.1 (Chen *et al*., 2018). Read adapters and poly-G tails at the end of the reads were trimmed. Filtered reads were assembled using MetaSPAdes v3.13.0 (Nurk *et al*., 2017). The Nuclear ribosomal RNA genes (5S, 5.8S, 18S, and 28S) were predicted from assemblies using Barrnap v0.9 (Seemann 2013). Mitochondrial genomes were retrieved and annotated using MitoFinder v1.4 (Allio *et al*., 2020). All genes were aligned separately using Mafft v7.305 (Katoh & Standley 2013). For the 13 mitochondrial protein-coding genes, we obtained using the transeq command of the EMBOSS v6.6.0 suite of programs (Rice *et al*., 2000) and carried out sequence alignment on the amino acid sequences. Amino acid sequence alignments were converted into DNA sequence alignments using PAL2NAL v14 (Suyama *et al*., 2006). Individual gene alignments were concatenated using FASconCAT-G (Kück & Longo 2014).The 22 mitochondrial transfer RNA genes, two ribosomal RNA genes (12S and 16S), and the six ribosomal RNA genes (mitochondrial 12S and 16S and nuclear 5S, 5.8S, 18S, and 28S) were aligned as DNA sequences, separately.

### 2.3 Phylogenetic analyses

All phylogenetic analyses were performed with and without the third codon positions of protein-coding genes. We reconstructed Maximum Likelihood phylogenetic trees using IQ-TREE 1.6.12 (Minh *et al*., 2020). The best-fit nucleotide substitution model was determined with ModelFinder (Kalyaanamoorthy *et al*., 2017) implemented in IQ-TREE v1.6.12. Branch supports were calculated using 1,000 bootstrap replicates (Hoang *et al*., 2018). Bayesian phylogenetic trees were inferred with MrBayes v3.2.3 using the GTR+G model of nucleotide substitution (Ronquist *et al*., 2012). The Markov chain Monte Carlo (MCMC) chains were run for 20 million generations for the datasets with or without the third codon positions of protein-coding genes. In all analyses, the MCMC chains were sampled every 5,000 generations to estimate the posterior distribution. The first 10% of sampled trees were excluded as burn-in. Visual inspection of the trace files with Tracer v1.7.1 confirmed that all analyses converged (Rambaut *et al*., 2018). The effective sample size was higher than 220 for every parameter of every run. The MCMC chains were run four times in parallel for both datasets.

### 2.4 Divergence time estimation

We reconstructed Bayesian time-calibrated phylogenies using BEAST v2.6.2 (Bouckaert *et al*., 2019). Bayesian analyses were performed with and without the third codon positions of protein-coding genes. We used an uncorrelated lognormal relaxed clock to model rate variation among branches. A Yule model was used as tree prior. A GTR+G model of nucleotide substitution was applied to each partition. The MCMC analyses were run for 100 million and 150 million generations for the analyses without and with third codon positions, respectively. The chains were sampled every 5,000 generations. We checked the convergence of the MCMC runs with Tracer v1.7.1 and consequently discarded the first 10% of generations as burn-in. We used ten fossils as time constraints (Table S2). Each calibration was implemented as an exponential prior on node time. The use of these calibrations has been thoroughly justified previously (Bucek *et al*., 2019, 2021). We used TreeAnnotator implemented in the BEAST2 suite of programs to generate a consensus tree. Tree topology and 95% height posterior density (HPD) were visualized with FigTree v 1.4.4 (Rambaut 2018).

## 3 Results

### 3.1 Phylogenetic reconstructions

The phylogenetic trees obtained using Maximum Likelihood and Bayesian analyses received high nodal support values and possessed almost identical topologies. One exception was the relationships among *Stolotermes inopinus* and the two samples of *Stolotermes ruficeps* that were resolved with low bootstrap values (<75%) and Bayesian posterior probabilities (<0.9) (Fig. 1).

**Fig. 1.**
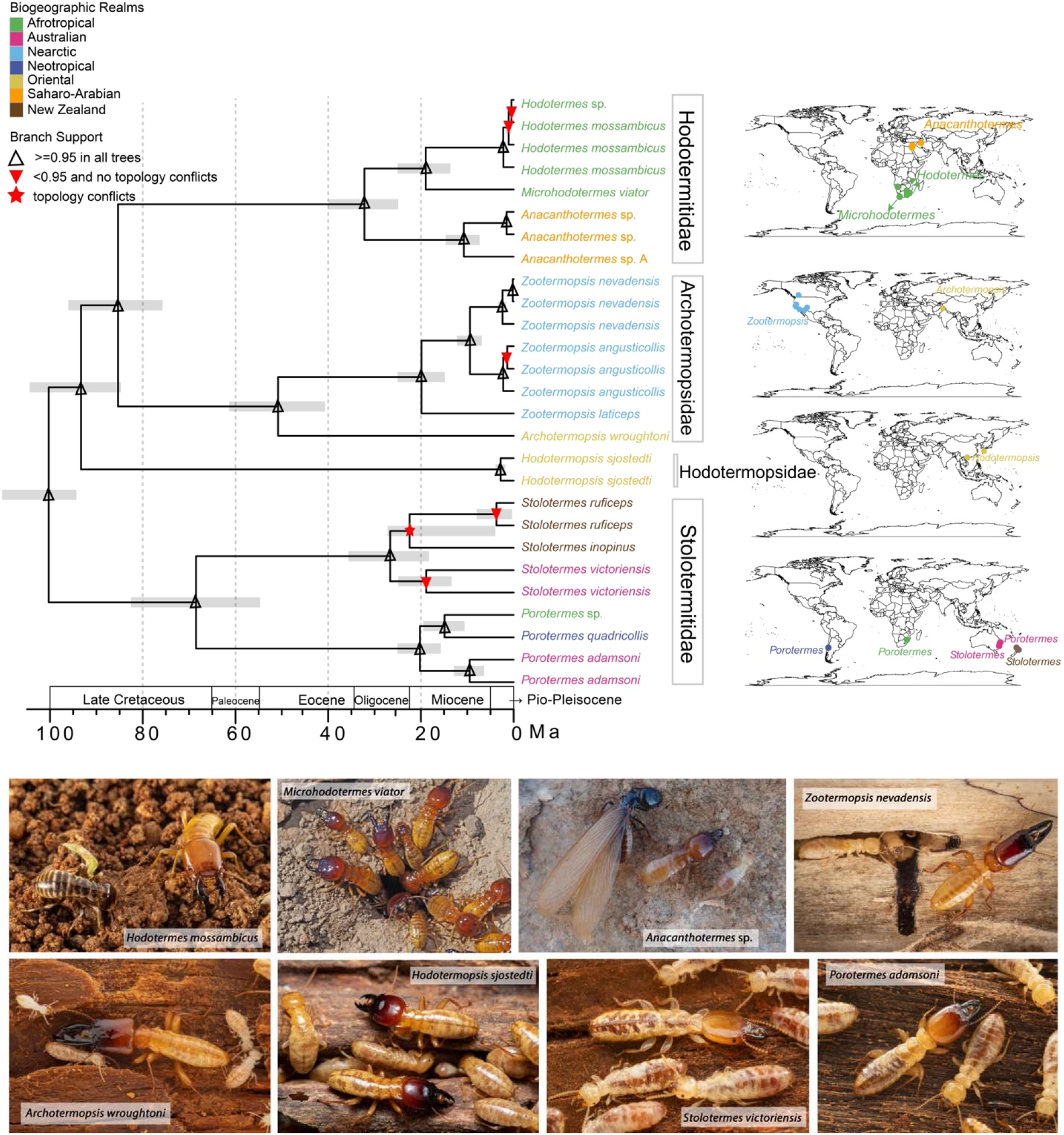
Time-calibrated phylogenetic tree of Stolotermitidae, Archotermopsidae, Hodotermopsidae, and Hodotermitidae based on full mitochondrial genomes and 5S, 5.8S, 18S, and 28S rRNA genes. The tree was reconstructed without third codon positions of protein-coding genes with BEAST2. The map shows the sampling locations of Stolotermitidae, Archotermopsidae, Hodotermopsidae, and Hodotermitidae. Node symbols (blank triangle, red triangle, and red stars) represent the bootstrap support and posterior probability values obtained with IQTREE, MrBayes, and BEAST2 on the dataset with and without third codon positions of protein-coding genes. Node bars indicate 95% Height Posterior Density intervals of age estimates. Biogeographic realms are given and based on the descriptions in Holt et al. 2013. Tip colors coincide with collect localities. Maps on the right show the collection localities of Hodotermitidae, Archotermopsidae, Hodotermopsidae, and Stolotermitidae. The photographs depict one species of each genus included in this study. Photographs of *Microhodotermes* and *Anacanthotermes* were provided by Felix Riegel and Omer Theodore, respectively.

Our analyses retrieved Mastotermitidae as sister group to Euisoptera, which comprised all non-matotermitid termites, and confirmed the monophyly of Stolotermitidae + Archotermopsidae + Hodotermitidae (Teletisoptera), which was retrieved as the sister group of Kalotermitidae + Neoisoptera (Icoisoptera). Stolotermitidae was found to be monophyletic and formed the sister group of Archotermopsidae + Hodotermitidae. The Archotermopsidae were retrieved as paraphyletic with respect to a monophyletic Hodotermitidae. Within the lineage composed of Archotermopsidae and Hodotermitidae, *Hodotermopsis* (Hodotermopsinae) was sister to the other five genera. *Zootermopsis* and *Archotermopsis* formed a monophyletic group sister to the three genera of Hodotermitidae (*i*.*e*., Archotermopsidae would be monophyletic with the removal of Hodotermopsinae). Within the Hodotermitidae, *Anacanthotermes* was found to be sister to *Hodotermes* + *Microhodotermes*. Each of the eight genera studied here were monophyletic.

### 3.2 Divergence dating

Time-calibrated phylogenies reconstructed with and without the third codon positions of protein-coding genes yielded similar time estimates, differing by less than three million years for each node. For this reason, we only provide the results of the analysis with the third codon position excluded (Fig. 1). The clade composed of Stolotermitidae, Hodotermitidae, and Archotermopsidae diverged from other Euisoptera 123.4 Ma (113.3–136.4 Ma, 95% HPD). Stolotermitidae diverged from Hodotermitidae + Archotermopsidae 100.3 Ma (94.3–110.4 Ma, 95% HPD). The most recent common ancestor of Stolotermitidae occurred around 68.5 Ma (54.7–82.5 Ma, 95% HPD), and the most recent common ancestors of *Porotermes* and *Stolotermes* were estimated to have existed 20.2 Ma (15.7–25.1 Ma, 95% HPD) and 26.6 Ma (18.3–35.6 Ma, 95% HPD), respectively. *Hodotermopsis* and other Archotermopsidae + Hodotermitidae diverged 93.3 Ma (84.7–104.4 Ma, 95% HPD). The divergence time of *Zootermopsis* and *Archotermopsis* was estimated to have occurred 50.8 Ma (40.7–61.4 Ma, 95% HPD), and the most recent common ancestor of *Zootermopsis* was estimated at 19.9 Ma (14.8–25.0 Ma, 95% HPD). Hodotermitidae diverged from *Zootermopsis* + *Archotermopsis* 85.3 Ma (75.7–96.1 Ma, 95% HPD). Within the Hodotermitidae, *Hodotermes* + *Microhodotermes* diverged from *Anacanthotermes* 32.2 Ma (24.9–40.0 Ma, 95% HPD). *Hodotermes* and *Microhodotermes* split 19.0 Ma (13.6–25.1 Ma, 95% HPD).

## 4 Discussion

In this study, we present a comprehensive phylogenetic reconstruction of the early diverging termite families Stolotermitidae, Archotermopsidae, and Hodotermitidae (Figs 1, S1). We used three phylogenetic reconstruction methods and repeated the analyses on datasets with and without third codon positions of protein-coding genes. The topology of the phylogenetic trees was largely consistent across methods and datasets, except for the positions of *Stolotermes inopinus* and *S. ruficeps*, which were discordant among analyses, probably owing to the limited amount of sequence data available for these two species. Our phylogenies were also congruent with previous estimates based on mitochondrial genomes and transcriptomes (Cameron *et al*., 2012; Bourguignon *et al*., 2015; Bucek *et al*., 2019). *Mastotermes* was found to be the sister group of Euisoptera, a clade comprising all other termites (Engel *et al*., 2009), and the group Stolotermitidae + Archotermopsidae + Hodotermitidae (Teletisoptera) was sister to Kalotermitidae + Neoisoptera (Icoisoptera). Our analyses supported the monophyly of Stolotermitidae, which was found to be sister Archotermopsidae + Hodotermidae, the former paraphyletic to the latter. The paraphyly of Archotermopsidae was already indicated by previous phylogenies based on full mitochondrial genomes (Bourguignon *et al*., 2015). It is clear that a simple augmentation of the current classification by removing *Hodotermopsis* from Archotermopsidae resolves this paraphyly, while simultaneously maximizing nomenclatural stability with the literature of the last 70 years (*i*.*e*., maintaining Grassé’s distinction between a family of harvesters and dampwood termites). Accordingly, we restrict Archotermopsidae to *Archotermopsis* and *Zootermopsis* (*i*.*e*., Archotermopsinae sensu Jiang *et al*. (2021) elevated as Archotermopsidae Engel et al., ***status novum***), and elevate Hodotermopsinae to familial rank (*i*.*e*., Hodotermopsidae Engel, ***status novum***). While this system is finely split, it is preferable to obscuring the biological differences and confusing the historical literature that has deployed these names, particularly Hodotermitidae, in such a context since Grassé (1949). The alteratives would be 1) recognizing all of the aforementioned families as subfamilies of Hodotermitidae (semantically equivalent to the multi-family system), or 2) to recognize two families, Stolotermitidae and Hodotermitidae, the former with Stolotermitinae and Porotermitinae, and the latter with Hodotermitinae, Archotermopsinae, and Hodotermopsinae. Neither of these alternatives maximize nomenclatural stability in the sense of the ICZN (1999), nor do they provide any greater clarity regarding relationships. Accordingly, the system we adopt (Table 1) emphasizes the ecological differences between the taxonomic units, with all Archotermopsidae and Hodotermopsidae feeding on damp wood (usually coniferous), while all Hodotermitidae are desert harvester termites feeding predominantly on dry grasses (Krishna *et al*., 2013). In the remainder of the discussion we shall refer to the families in this new context.

The time-calibrated trees estimated with and without third codon positions of protein-coding genes yielded similar time estimates. Our time estimates of the branching among early diverging termite families were younger than those of previous studies. However, our estimates were largely congruent with overlapping HPD intervals. For example, we estimated the most recent common ancestor of termites at 133.1 Ma (125.5–145.4 Ma, 95% HPD), while previous studies found older ages: 149 Ma (136–170 Ma, 95% HPD) (Bourguignon *et al*., 2015), 151.3 Ma(149.3–153.7 Ma, 95% HPD) (Legendre *et al*., 2015), and 140.6 Ma (112.6–170.5 Ma, 95% HPD) (Bucek *et al*., 2019). Differences among studies in terms of fossil calibrations, fossil age estimations, taxonomic sampling, and models used for the reconstruction of time-calibrated trees may be the causes of this variation.

We did not attempt to reconstruct the ancestral range of Stolotermitidae + Hodotermopsidae + Archotermopsidae + Hodotermitidae, particularly given that the many fossils occurring well outside of modern distributions would render meaningless such an estimate based solely on extant taxa. Ancestral range reconstructions have been performed previously for Neoisoptera and Kalotermitidae (Bourguignon *et al*., 2016, 2017; Wang *et al*., 2019; Romero Arias *et al*., 2021; Bucek *et al*., 2021). However, compared to Stolotermitidae + Hodotermopsidae + Archotermopsidae + Hodotermitidae, Neoisoptera and Kalotermitidae are diverse and widespread, comprising many extant species whose distribution and phylogenetic relationships can inform on past vicariance and dispersal events, and with most fossils nested within those distributions (Krishna *et al*., 2013). Stolotermitidae, Hodotermopsidae, Archotermopsidae, and Hodotermitidae are species-poor families, with limited modern distributions, relict of past wider distributions as evidenced from the fossil record (Krishna *et al*., 2013; Engel *et al*., 2013, 2016; Jiang *et al*., 2021).Most geographic lineages of Teletisoptera inhabit regions hosting few other termites and may have been competitively excluded from regions where termitids and other Neoisoptera became dominant during the Oligocene and Miocene (Engel *et al*., 2009; Bourguignon *et al*.,2017). Teletisoptera inhabit regions generally devoid of other members of the group, preventing a meaningful reconstruction of its historical biogeography.

While the low diversity of teletisopteran families hamper meaningful ancestral range reconstructions, our time-calibrated trees permit the identification of several biogeographic disjunctions. The two modern stolotermitid genera, *Porotermes* and *Stolotermes*, have a Gondwanan distribution (Emerson 1942, 1955; Gay & Calaby 1969; Kaulfuss *et al*., 2010; Krishna *et al*., 2013). However, our time-calibrated phylogeny indicated that all species of *Porotermes* share a common ancestor 20.2 Ma (15.7–25.1 Ma, 95% HPD) and the common ancestor of the species of *Stolotermes* sequenced in this study lived 26.6 Ma (18.3–35.6 Ma, 95% HPD), both considerably younger than the breakup of Gondwana. Although we could not sequence *S. africanus*, the only species of *Stolotermes* found in Africa, our time-calibrated trees showed that *Stolotermes* diverged from *Porotermes* 68.5 Ma (54.7–82.3 Ma, 95% HPD), after the breakup of Gondwana. Interestingly, an extinct genus allied to *Porotermes* is known from the Oligocene of Ethiopia (Engel *et al*., 2013), predating the divergence of crown-group *Porotermes* but postdating the divergence of the lineages comprising Porotermitinae and Solotermitinae. Collectively, these results imply that the presence of *Stolotermes* in South Africa, eastern Australia as well as New Zealand, and the presence of *Porotermes* in southern Australia, southern Africa, and South America is not the result of vicariance during the breakup of Gondwana, as hypothesized previously (Krishna *et al*., 2013; Bourguignon *et al*., 2015). Instead, *Porotermes* and *Stolotermes* acquired their modern distribution through long-distance oversea dispersal events.

The biogeographic disjunctions among modern genera of Hodotermopsidae + Archotermopsidae + Hodotermitidae may be explained by land bridges. Indeed, we estimated that Hodotermopsidae + Archotermopsidae + Hodotermitidae shared a common ancestor around 93.3 Ma (84.7–104.4 Ma, 95% HPD), indicating vicariance through continental drift may explain the distribution of early diverging members of this clade. The Palearctic region remained connected to North America through Greenland until about 50 Ma (Scotese 2004), possibly explaining the disjunction between the Palearctic *Archotermopsis* and the Nearctic *Zootermopsis*, the modern descendants of more widespread ancestors (Krishna *et al*., 2013). The African *Hodotermes* + *Microhodotermes* diverged from *Anacanthotermes*, a genus found in Africa, the Middle East, and South Asia, 32.2 Ma (24.9–40.0 Ma, 95% HPD) and the most recent common ancestors of *Hodotermes* + *Microhodotermes* and *Anacanthotermes* lived 19.0 Ma (13.6–25.1 Ma, 95% HPD) and 10.7 Ma (7.3–14.7 Ma, 95% HPD), respectively. The timing of the biogeographic disjunction between these two lineages may coincide with the existence of the *Gomphotherium* land bridge that connected Africa and Eurasia ∼18–20 Ma (Rögl 1998, 1999). The sequencing of African *Anacanthotermes* in future studies is needed to confirm this scenario.

Our study showcases the importance of samples collected before the genomics era for future phylogenetic reconstructions. One limitation of many studies attempting to reconstruct the evolution of diverse taxa is the sampling of a representative set of specimens covering the diversity of the groups of interest. Because species of Stolotermitidae, Hodotermopsidae, Archotermopsidae, and Hodotermitidae occur in regions where termite diversity is generally low, we made fewer attempts to collect them. Instead, this study is largely based on samples collected in ethanol during the last three decades for taxonomic purposes. In addition, we sequenced a syntype of *Archotermopsis wroughtoni* (Desneux, 1904), that was collected in the Kashmir Valley. The systematic sequencing of type material, such as a syntype of *A. wroughtoni* sequenced in this study, holds the promise of clarifying the taxonomic literature and making available type-based species identification to the whole scientific community.

## Acknowledgments

Rudi Scheffrahn provided specimens and commented this manuscript. We also thank the DNA Sequencing Section and the Scientific Computation and Data Analysis Section of the Okinawa Institute of Science and Technology Graduate University, Okinawa, Japan, for assistance with sequencing and for providing access to the OIST computing cluster, respectively. This work was supported by the Japan Society for the Promotion of Science through a DC2 graduate student fellowship to JA and a postdoctoral fellowship to SH (19F19819). We also acknowledge support from the Internal Grant Agency of the Faculty of Tropical AgriSciences (CULS No. 20213112). We thank Felix Riegel and Omer Theodore for providing the photographs of *Microhodotermes* and *Anacanthotermes*, respectively.

## Notes

### Competing Interest Statement

The authors have declared no competing interest.

## References

Allio, R., Schomaker-Bastos, A., Romiguier, J., Prosdocimi, F., Nabholz, B. & Delsuc, F. (2020) MitoFinder: efficient automated large-scale extraction of mitogenomic data in target enrichment phylogenomics. Molecular Ecology Resources, 20, 892–905.

Barden, P. & Engel, M.S. (2021) Fossil Social Insects. In: Encyclopedia of Social Insects. pp. 384–403. Springer, Cham.

Barrett, J.H. (1965) The occurrence of termites in the New Guinea highlands. Papua and New Guinea Agricultural JournaL, 17, 95–98.

Bezerra, F.I., Mendes, M. & De Souza, O. (2020) New record of Mastotermitidae from Fonseca Basin, Eocene-Oligocene boundary of southeastern Brazil. Biologia, 75, 1881–1890.

Bouckaert, R., Vaughan, T.G., Barido-Sottani, J., Duchêne, S., Fourment, M., Gavryushkina, A., Heled, J., Jones, G., Kühnert, D., De Maio, N., Matschiner, M., Mendes, F.K., Müller, N.F., Ogilvie, H.A., Du Plessis, L., Popinga, A., Rambaut, A., Rasmussen, D., Siveroni, I., Suchard, M.A., Wu, C.H., Xie, D., Zhang, C., Stadler, T. & Drummond, A.J. (2019) BEAST 2.5: an advanced software platform for bayesian evolutionary analysis. PLoS Computational Biology, 15, e1006650.

Bourguignon, T., Lo, N., Cameron, S.L., Šobotník, J., Hayashi, Y., Shigenobu, S., Watanabe, D., Roisin, Y., Miura, T. & Evans, T.A. (2015) The evolutionary history of termites as inferred from 66 mitochondrial genomes. Molecular Biology and Evolution, 32, 406–421.

Bourguignon, T., Lo, N., Šobotník, J., Ho, S.Y.W., Iqbal, N., Coissac, E., Lee, M., Jendryka, M.M., Sillam-Dussès, D., Křížková, B., Roisin, Y. & Evans, T.A. (2017) Mitochondrial phylogenomics resolves the global spread of higher termites, ecosystem engineers of the tropics. Molecular Biology and Evolution, 34, 589–597.

Bourguignon, T., Lo, N., Šobotník, J., Sillam-Dussès, D., Roisin, Y. & Evans, T.A. (2016) Oceanic dispersal, vicariance and human introduction shaped the modern distribution of the termites Reticulitermes, Heterotermes and Coptotermes. Proceedings of the Royal Society B: Biological Sciences, 283, 20160179.

Bucek, A., Šobotník, J., He, S., Shi, M., McMahon, D.P., Holmes, E.C., Roisin, Y., Lo, N. & Bourguignon, T. (2019) Evolution of termite symbiosis informed by transcriptome-based phylogenies. Current Biology, 29, 3728–3734.

Bucek, A., Wang, M., Sobotník, J., Sillam-Dussès, D., Mizumoto, N., Stiblik, P., Clitheroe, C., Lu, T., Gonzalez, P.J.J., Mohagan, A., Rafanomezantsoa, J.J., Fisher, B., Engel, M.S., Roisin, Y., Evans, T.A., Scheffrahn, R.H. & Bourguignon, T. (2021) Transoceanic voyages of “drywood” termites (Isoptera: Kalotermitidae) inferred from extant and extinct species. bioRxiv, unpublished.

Cameron, S.L., Lo, N., Bourguignon, T., Svenson, G.J. & Evans, T.A. (2012) A mitochondrial genome phylogeny of termites (Blattodea: Termitoidae): robust support for interfamilial relationships and molecular synapomorphies define major clades. Molecular Phylogenetics and Evolution, 65, 163–173.

Chen, S., Zhou, Y., Chen, Y. & Gu, J. (2018) Fastp: an ultra-fast all-in-one FASTQ preprocessor. In: Bioinformatics. pp. i884–i890. Oxford University Press.

Edwards, E.J., Osborne, C.P., Stromberg, C.A.E., Smith, S.A., Bond, W.J., Christin, P.A., Cousins, A.B., Duvall, M.R., Fox, D.L., Freckleton, R.P., Ghannoum, O., Hartwell, J., Huang, Y., Janis, C.M., Keeley, J.E., Kellogg, E.A., Knapp, A.K., Leakey, A.D.B., Nelson, D.M., Saarela, J.M., Sage, R.F., Sala, O.E., Salamin, N., Still, C.J. & Tipple, B. (2010) The origins of C4 grasslands: integrating evolutionary and ecosystem science. Science, 328, 587–591.

Emerson, A.E. (1933) A revision of the genera of fossil and recent Termopsinae (Isoptera). University of California Press, Berkeley,.

Emerson, A.E. (1967) Cretaceous insects from Labrador 3. a new genus and species of termite (Isoptera: Hodotermitidae). Psyche, 74, 276–289.

Emerson, A.E. (1955) Geographical origins and dispersions of termite genera. Fieldiana: Zoology, 37, 465–521.

Emerson, A.E. (1942) The relations of a relict South African termite (Isoptera, Hodotermitidae, Stolotermes). American Museum Novitates, 1–12.

Engel, M.S., Barden, P., Riccio, M.L. & Grimaldi, D.A. (2016) Morphologically specialized termite castes and advanced sociality in the early Cretaceous. Current Biology, 26, 522–530.

Engel, M.S., Currano, E.D. & Jacobs, B.F. (2015) The first mastotermitid termite from Africa (Isoptera: Mastotermitidae): a new species of Mastotermes from the early Miocene of Ethiopia. Journal of Paleontology, 89, 1038–1042.

Engel, M.S., Grimaldi, D.A. & Krishna, K. (2007) A synopsis of Baltic amber termites (Isoptera). Stuttgarter Beiträge zur Naturkunde, Serie B (Geologie und Paläontologie), 372, 1–20.

Engel, M.S., Grimaldi, D.A. & Krishna, K. (2009) Termites (Isoptera): their phylogeny, classification, and rise to ecological dominance. American Museum Novitates, 2009, 1–27.

Engel, M.S. & Gross, M. (2009) A giant termite from the Late Miocene of Styria, Austria (Isoptera). Naturwissenschaften, 96, 289–295.

Engel, M.S. & Krishna, K. (2004) Family-group names for termites (Isoptera). American Museum Novitates, 3432, 1–9.

Engel, M.S., Pan, A.D. & Jacobs, B.F. (2013) A termite from the late Oligocene of Northern Ethiopia. Acta Palaeontologica Polonica, 58, 331–334.

Engel, M.S. & Tanaka, T. (2015) A giant termite of the genus Gyatermes from the late Miocene of Nagano Prefecture, Japan (Isoptera). Novitates Paleoentomologicae, 1–12.

Gay, F.J. & Calaby, J.H. (1969) Termites of the Australian regions. In: Biology of Termites. pp. 398–448. Springer-Verlag.

Grassé, P.P. (1949) Termites. Isoprères. Traité de Zoologie, 9, 408–544.

Grimaldi, D.A. & Engel, M.S. (2005) Evolution of the insects. Cambridge University Press.

Hoang, D.T., Chernomor, O., Von Haeseler, A., Minh, B.Q. & Vinh, L.S. (2018) UFBoot2: improving the ultrafast bootstrap approximation. Molecular Biology and Evolution, 35, 518–522.

Holmgren, N. (1911) Termiten studien 2. Systematic der Termiten. Die Familie Mastotermitidae, Protermitidae und Mesotermitidae. Kungliga Svenska Vetenskapsakademiens Handlingar. 46, 1–86.

Holt, B.G., Lessard, J.-P., Borregaard, M.K., Fritz, S.A., Araujo, M.B., Dimitrov, D., Fabre, P.-H., Graham, C.H., Graves, G.R., Jonsson, K.A., Nogues-Bravo, D., Wang, Z., Whittaker, R.J., Fjeldsa, J. & Rahbek, C. (2013) An update of Wallace’s zoogeographic regions of the world. Science, 339, 74–78.

Inward, D., Beccaloni, G. & Eggleton, P. (2007a) Death of an order: a comprehensive molecular phylogenetic study confirms that termites are eusocial cockroaches. Biology Letters, 3, 331–335.

Inward, D.J.G., Vogler, A.P. & Eggleton, P. (2007b) A comprehensive phylogenetic analysis of termites (Isoptera) illuminates key aspects of their evolutionary biology. Molecular Phylogenetics and Evolution, 44, 953–967.

Jiang, R., Zhang, H., Eldredge, K.T., Song, X., Li, Y., Tihelka, E., Huang, D., Wang, S., Engel, S. M., & Cai, C. (2021) Further evidence of Cretaceous termitophily: description of new termite hosts of the trichopseniine Cretotrichopsenius (Coleoptera: Staphylinidae), with emendations to the classification of lower termites (Isoptera). Palaeoentomology, 4, 374–389.

Kalyaanamoorthy, S., Minh, B.Q., Wong, T.K.F., Von Haeseler, A. & Jermiin, L.S. (2017) ModelFinder: fast model selection for accurate phylogenetic estimates. Nature Methods, 14, 587–589.

Katoh, K. & Standley, D.M. (2013) MAFFT multiple sequence alignment software version 7: improvements in performance and usability. Molecular Biology and Evolution, 30, 772–780.

Kaulfuss, U., Harris, A.C. & Lee, D.E. (2010) A new fossil termite (Isoptera, Stolotermitidae, Stolotermes) from the Early Miocene of Otago, New Zealand. Acta Geologica Sinica - English Edition, 84, 705–709.

Krishna, K. (1970) Taxonomy, phylogeny, and distribution of termites. Biology of Termites, 127–152.

Krishna, K. & Emerson, A.E. (1983) A new fossil species of termite from Mexican amber, Mastotermes electromexicus (Isoptera, Mastotermitidae). American Museum of Natural History, 2767, 1–8.

Krishna, K. & Grimaldi, D. (1991) A new fossil species from Dominican amber of the living Australian termite genus Mastotermes (Isoptera: Mastotermitidae). American Museum of Natural History, 1–10.

Krishna, K. & Grimaldi, D.A. (2000) A new subfamily, genus, and species of termite (Isoptera) from New Jersey Cretaceous amber. Grimaldi, David[Ed]. In: Studies on fossils in amber, with particular reference to the Cretaceous of New Jersey. pp. 133–140. Backhuys Publishers, Leiden, The Netherlands.

Krishna, K., Grimaldi, D.A., Krishna, V. & Engel, M.S. (2013) Treatise on the Isoptera of the world. Bulletin of the American Museum of Natural History, 377, 1–200.

Kück, P. & Longo, G.C. (2014) FASconCAT-G: extensive functions for multiple sequence alignment preparations concerning phylogenetic studies. Frontiers in Zoology, 11, 81.

Legendre, F., Nel, A., Svenson, G.J., Robillard, T., Pellens, R. & Grandcolas, P. (2015) Phylogeny of Dictyoptera: dating the origin of cockroaches, praying mantises and termites with molecular data and controlled fossil evidence. PLoS ONE, 10, e0130127.

Lo, N., Tokuda, G., Watanabe, H., Rose, H., Slaytor, M., Maekawa, K., Bandi, C. & Noda, H. (2000) Evidence from multiple gene seqeunces indicates that termites evolved from wood-feeding cockroaches. Current Biology, 10, 801–804.

Minh, B.Q., Schmidt, H.A., Chernomor, O., Schrempf, D., Woodhams, M.D., Von Haeseler, A., Lanfear, R. & Teeling, E. (2020) IQ-TREE 2: new models and efficient methods for phylogenetic inference in the genomic era. Molecular Biology and Evolution, 37, 1530–1534.

Nurk, S., Meleshko, D., Korobeynikov, A. & Pevzner, P.A. (2017) MetaSPAdes: a new versatile metagenomic assembler. Genome Research, 27, 824–834.

Rambaut, A. (2018) FigTree v1.4.4. http://tree.bio.ed.ac.uk/software/figtree/,.

Rambaut, A., Drummond, A.J., Xie, D., Baele, G. & Suchard, M.A. (2018) Posterior summarization in bayesian phylogenetics using Tracer 1.7. Systematic Biology, 67, 901–904.

Rice, P., Longden, I. & Bleasby, A. (2000) EMBOSS: the European molecular biology open software suite. Trends in Genetics, 16, 276–277.

Rögl, F. (1999) Mediterranean and Paratethys. Facts and hypothesis of an Oligocene to Miocene paleogeography (short review). Annalen des Naturhistorischen Museums in Wien, A1014, 339–349.

Rögl, F. (1998) Palaeogeographic considerations for Mediterranean and Paratethys Seaways (Oligocene to Miocene). Annalen des Naturhistorischen Museums in Wien, 99A, 279–310.

Romero Arias, J., Boom, A., Wang, M., Clitheroe, C., Šobotník, J., Stiblik, P., Bourguignon, T. & Roisin, Y. (2021) Molecular phylogeny and historical biogeography of Apicotermitinae (Blattodea: Termitidae). Systematic Entomology, 46, 741–756.

Ronquist, F., Teslenko, M., van der Mark, P., Ayres, D.L., Darling, A., Höhna, S., Larget, B., Liu, L., Suchard, M.A. & Huelsenbeck, J.P. (2012) MrBayes 3.2: efficient bayesian phylogenetic Inference and model choice across a large model space. Systematic Biology, 61, 539–542.

Scotese, R.S. (2004) Cenozoic and Mesozoic paleogeography: changing terrestrial biogeographic pathways. In: Frontiers of Biogeography: New directions in the geography of nature. (Ed. M.V.L. and L.R. Heaney), pp. 9–26. Sunderland, Massachusetts.

Seemann, T. (2013) barrnap 0.9LJ: rapid ribosomal RNA prediction. Github.com,.

Snyder, E.A.E. (1949) Catalog of the termites (Isoptera) of the world. Smithsonian miscellaneous collections, 112, 1–490.

Suyama, M., Torrents, D. & Bork, P. (2006) PAL2NAL: robust conversion of protein sequence alignments into the corresponding codon alignments. Nucleic Acids Research, 34, W609–W612.

Thorne, B.L., Grimaldi, D.A. & Krishna, K. (2000) Early fossil history of the termites. In: Termites: Evolution, Sociality, Symbioses, Ecology. pp. 77–93. Springer Netherlands.

Vršanský, P. & Aristov, D. (2014) Termites (Isoptera) from the Jurassic/Cretaceous boundary: evidence for the longevity of their earliest genera. European Journal of Entomology, 111, 137–141.

Wang, M., Buček, A., Šobotník, J., Sillam-Dussès, D., Evans, T.A., Roisin, Y., Lo, N. & Bourguignon, T. (2019) Historical biogeography of the termite clade Rhinotermitinae (Blattodea: Isoptera). Molecular Phylogenetics and Evolution, 132, 100–104.

Wappler, T. & Engel, M.S. (2006) A new record of Mastotermes from the Eocene of Germany (Isoptera: Mastotertitidae). Journal of Paleontology, 80, 380–385.

Weidner, H. (1955) Körperbau, Systematik und Verbreitung der Termiten. In Schmidt, H., Die Termiten. Leipzig, Akad. Verlagsgesellschaft, Geest und Portig. 5–81.

Yashiro, T., Mitaka, Y., Nozaki, T. & Matsuura, K. (2018) Chemical and molecular identification of the invasive termite Zootermopsis nevadensis (Isoptera: Archotermopsidae) in Japan. Applied Entomology and Zoology, 53, 215–221.

Zachos, J., Pagani, H., Sloan, L., Thomas, E. & Billups, K. (2001) Trends, rhythms, and aberrations in global climate 65 Ma to present. Science, 292, 686–693.

Zhao, Z., Eggleton, P., Yin, X., Gao, T., Shih, C. & Ren, D. (2019) The oldest known mastotermitids (Blattodea: Termitoidae) and phylogeny of basal termites. Systematic Entomology, 44, 612–623.

