## Supplementary material for "Historical biogeography of early diverging termite lineages (Isoptera: Teletisoptera)"

**Fig. S1.** Time-calibrated phylogenetic tree of 64 termite samples reconstructed with BEAST2 using mitochondrial genomes without third codon positions and 5S, 5.8S, 18S, and 28S rRNA genes. Node bars indicate the 95% Height Posterior Density intervals. Branch labels represent posterior probabilities.

**Table S1.** Samples used in this study with corresponding collection details and accession numbers.

**Table S2.** Fossils used for time calibration in this study.

**Supplementary references**

Emerson, A.E. (1971) Tertiary fossil species of the Rhinotermitidae (Isoptera), phylogeny of genera, and reciprocal phylogeny of associated Flagellata (Protozoa) and the Staphylinidae (Coleoptera). *Bulletin of the American Museum of Natural History*, **146**, 245–303.

Engel, M.S., Grimald, D.A., Nascimbene, P.C. & Singh, H. (2011) The termites of Early Eocene Cambay amber, with the earliest record of the Termitidae (Isoptera). *ZooKeys*, **148**, 105–123.

Engel, M.S., Grimaldi, D.A. & Krishna, K. (2007a) A synopsis of Baltic amber termites (Isoptera). *Stuttgarter Beiträge zur Naturkunde, Serie B (Geologie und Paläontologie)*, **372**, 1–20.

Engel, M.S., Grimaldi, D.A. & Krishna, K. (2007b) Primitive termites from the Early Cretaceous of Asia (Isoptera). *Stuttgarter Beiträge zur Naturkunde, Serie B (Geologie und Paläontologie)*, **371**, 1–32.

Krishna, K. (1996) New fossil species of termites of the subfamily Nasutitermitinae from Dominican and Mexican amber (Isoptera, Termitidae). *American Museum Novitates*, **3176**, 1–13.

Krishna, K. & Grimaldi, D. (2009) Diverse Rhinotermitidae and Termitidae (Isoptera) in Dominican Amber. *American Museum Novitates*, **2009**, 1–48.

Krishna, K. & Grimaldi, D.A. (2003) The first Cretaceous Rhinotermitidae (Isoptera): a new species, genus, and subfamily in Burmese amber. *American Museum Novitates*, **3390**, 1–10.

Martins-Neto, R.G., Mancuso, A. & Gallego, O.F. (2005) The Triassic insect fauna from Argentina. Blattoptera from the Los Rastros Formation (Bermejo Basin), La Rioja Province. *Ameghiniana*, **42**, 705–723.

Schlemmermeyer, T. & Cancello, E.M. (2000) New fossil termite species: *Dolichorhinotermes dominicanus* from Dominican amber (Isoptera, Rhinotermitidae, Rhinotermitinae). *Papeis Avulsos de Zoologia*, **41**, 303–311.

Zhao, Z., Yin, X., Shih, C., Gao, T. & Ren, D. (2019) Termite colonies from mid-Cretaceous Myanmar demonstrate their early eusocial lifestyle in damp wood. *National Science Review*, **7**, 381–390.
