## Supplementary figures and images for "Historical biogeography of early diverging termite lineages (Isoptera: Teletisoptera)"

### Figure S1

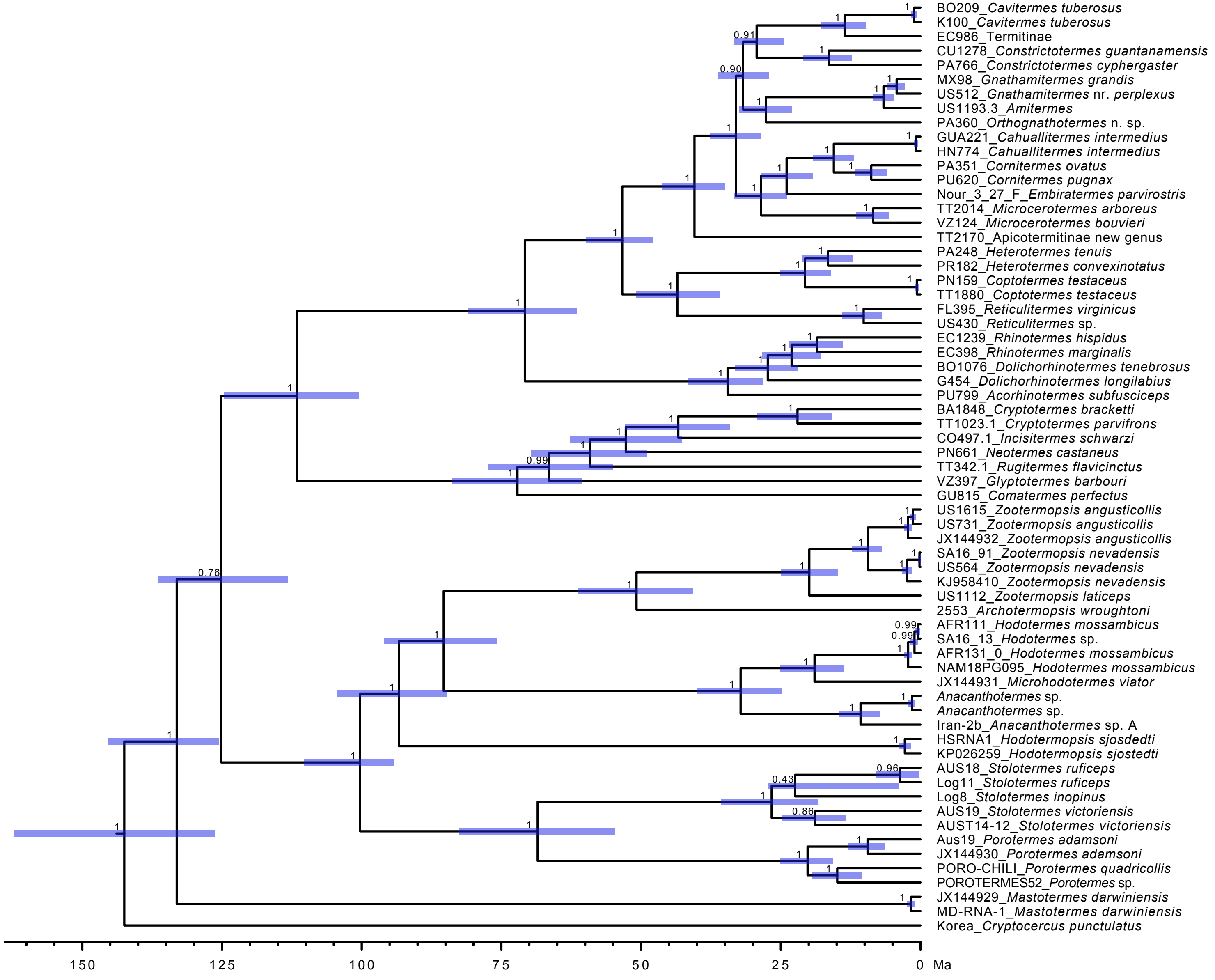
